# Sex, Competition and Mimicry : an eco-evolutionary model reveals how ecological interactions shape the evolution of phenotypes in sympatry

**DOI:** 10.1101/2020.11.30.403410

**Authors:** Boussens-Dumon Grégoire, Llaurens Violaine

**Affiliations:** Institut de Systématique, Evolution et Biodiversité (UMR 7205 CNRS/MNHN/SU/EPHE/UA), Muséum National d’Histoire Naturelle - CP50, 57 rue Cuvier 75005 PARIS, FRANCE

**Keywords:** mate preference, species recognition, mimetic coloration, divergence, density-dependent selection, mutualism

## Abstract

Phenotypic evolution in sympatric species can be strongly impacted by species interactions, either mutualistic or antagonistic, which may favour local phenotypic divergence or convergence. Interspecific sexual interactions between sympatric species has been shown to favour phenotypic divergence of traits used as sexual cues for example. Those traits may also be involved in local adaptation or in other types of species interactions resulting in complex evolution of traits shared by sympatric species. Here we focus on mimicry and study how reproductive interference may impair phenotypic convergence between species with various levels of defences. We use a deterministic model assuming two sympatric species and where individuals can display two different warning colour patterns. This eco-evolutionary model explores how ecological interactions shape phenotypic evolution within sympatric species. We investigate the effect of (1) the opposing density-dependent selections exerted on colour patterns by predation and reproductive behaviour, and (2) the impact of relative species and phenotype abundances on the fitness costs faced by each individual depending on their species and phenotype. Our model shows that reproductive interference may limit the convergent effect of mimetic interactions and may promote phenotypic divergence between Müllerian mimics. The divergent and convergent evolution of traits also strongly depends on the relative species and phenotype abundances and levels of trophic competition, highlighting how the eco-evolutionary feedbacks between phenotypic evolution and species abundances may result in strikingly different evolutionary routes.

## 2 Introduction

Phenotypic diversification among species is mainly influenced by neutral divergence, driven by the lack of gene flow between species. However, ecological interactions, either antagonistic or mutualistic play a significant role in the evolution of certain traits in sympatric species [6, 7, 28]. A striking example of mutualistic interactions shaping trait convergence in sympatric species is Müllerian mimicry. Predators attacking defended prey (e.g. prey containing deterrent chemical compounds) that display a warning signal (e.g. bright coloration) can learn to associate that warning signal with the defensive traits [27, 30]. This results in density-dependent selection because predation on defended individuals sharing the same warning signal decreases when their number increases [9, 26]. This favours local convergence towards the most commonly displayed warning signal across sympatric species with similar levels of defences [5, 16]. Such convergence of sympatric defended species’ phenotypes towards a similar warning signal has been documented in various taxa, including notably butterflies [5] and poison-frogs [30], forming so-called mimicry rings composed of multiple species sharing similar warning coloration and micro-habitat [7].

Despite the strong positive density-dependent selection exerted by predators, different warning signals are often maintained in sympatry [3, 15]. Although distribution of the different signals in different micro-habitats might favour such coexistence [4, 10], this puzzling observation is still largely unexplained. On top of contrasted developmental constraints in distantly-related species, founder effect and other stochastic factors affecting population demography might promote divergence among localities [22], but the factors determining the diversification of warning patterns among sympatric species involved in Müllerian mimicry are still largely unknown.

A poorly explored hypothesis relies on the implication of warning colour pattern in mate recognition. Warning colour patterns are indeed often involved in mate detection and selection [25]. When colour-pattern based assortative mating is predominant, increased resemblance between heterospecific sympatric individuals can result in increased sexual interactions across species. These interactions range from signal jamming (conspecific signals degraded by other species signals) to heterospecific courtship and ultimately hybridization. In the case of low-fitness hybrids or complete reproductive isolation of the sympatric species, these reproductive interferences (RI) can be very costly in time and energy (see [12] for a review). The cost of RI can then result in a segregation of spatial or temporal niches [19, 31], character displacement [21, 18] and sometimes even in species extinction [28]. The strength of RI depends on the encountering rates between males and females of sympatric species, and thus on the relative species density and niche overlap [31, 33]. Because mimicry and reproductive interference are both density-dependent, the local densities of each species and each phenotype has thus a crucial impact on the intensity of the evolutionary pressures faced by each individual.

Here we explore how pressures resulting from RI impact phenotypic changes within each species and especially how it may limit the evolutionary convergence in colour patterns due to mimicry between sympatric defended species. We use a differential equation model assuming two sympatric species displaying colour pattern variations, taking into account (1) the positive density-dependent selection on colour patterns across species due to predator behaviour, and (2) the relative species abundances that may favour the evolution of mimetic colour pattern in these two species. We explore how the benefit of mimicry, the cost of RI, and the strength of mate preferences on colour pattern may influence the evolution of colour pattern in sympatric species.

## 3 Materials and Methods

### 3.1 Purpose of the model

We aim at understanding how ecological interactions shape the evolution of a trait shared by two sympatric species. We particularly focus on the eco-evolutionary feedbacks, whereby trait variations within species and species densities influence each other. We perform simulations of the density-dependent evolutionary processes involved in colour pattern variations within and between two sympatric species, focusing on selective pressures generated by phenotypic preferences, reproductive interference and mimicry. We explicitly simulate the evolution of the densities of two different phenotypes within two sympatric species, resulting in a 4-equations system. We investigate conditions enabling either phenotypic convergence or divergence between the two sympatric species.

### 3.2 Model assumptions

We use deterministic simulations assuming no genetic drift and no mutations to investigate the evolution in the number of individuals displaying a given phenotype within each species. The two different phenotypes considered for each species are labelled A and B. The modelling of the different evolutionary pressures follow the assumptions listed below.

- <u>“Perfect” mimicry</u>. We assume a “perfect” mimicry, *i.e.* predators cannot differentiate two mimetic individuals, even if they belong to different species. As a result, **conspecific and heterospecific individuals with the same colour pattern both contribute to predator learning for that pattern**. The extent of their contribution only depends on their respective abundances and defence levels **L**, mimicry being density-dependent (see eq. 4).
- <u>Assortative mating</u>. We assume two haploid species with a single locus controlling the colour pattern variation. One allele codes for the phenotype A and one for the phenotype B. **Individuals can mate with any sympatric individual but we consider they do so more often with individuals sharing the same phenotype** (i.e. assortative mating) as often observed in mimetic species. Their tendency to mate with individuals of the alternative phenotype is given by **P** ∈ [0, 1] (see eq. 3). Because both species share the same phenotypes A and B, individuals may also engage in reproductive behaviours with mimetic individual from the alternative species, resulting in reproductive interference.
- <u>Reproductive interference (RI)</u>. We consider a strict post-zygotic isolation, so that crosses between individuals belonging to different species leave no offspring. However, **individuals can still mistakenly court heterospecific individuals, because of their shared phenotype and tendency to assortative mating**. The lack of viable offspring resulting from these heterospecific sexual interactions generates a reproductive interference cost. This cost depends on individuals tendency to engage in heterospecific reproductive behaviours, given by **I** ∈[0, 1] (see eq. 3) and colour pattern preferences, given by **P**.
- <u>Trophic competition</u>. Competition for resource occurs within and between the two species. The intensity of the competition for resource is assumed to be proportional to species density and to the degree of niche overlap, described by *c*. We consider a partial niche overlap between species, *c*_*b*_ < 1, and a complete niche overlap within species, *c*_*w*_ = 1 (see eq. 2).

### 3.3 Life cycle

The number of individuals of each phenotype within each species can change through time depending on the rate of generation of offsprings (*G*) and on adult survival (*S*), as described in equation 1 for individuals of species 1 and phenotype A.

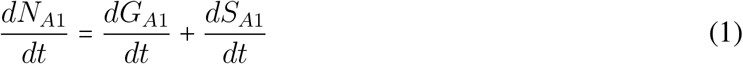

We consider all death, birth and mating events simultaneously. Equations 2 to 5 detail the offspring generation and adult survival terms. For clarity purposes, we focuse on the number of individuals of species 1 displaying the phenotype A, but the detailed equations for the four individuals types can be found in supplementary material (S1.1).

#### 3.3.1 Offspring generation

The production of individuals of a given phenotype within each species are based on Lotka-Volterra competition equations, taking into account both reproductive success and offspring survival. In sympatric species, resource limitation may indeed strongly affect survival in early developmental stages : in mimetic butterflies for instance, larval competition within and among species for host-plants is known to shape species co-existence [32]. The contribution of offspring generation *G*_*A*1_ to the change in number of individuals with phenotype A in species 1 is thus modelled by :

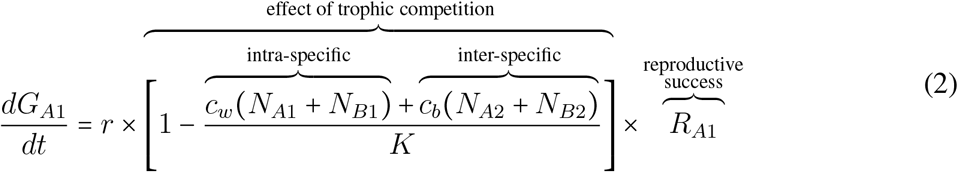

with

- *r* : the basal birth rate, equal for all individuals regardless of species or phenotype.
- *c*_*w*_ : the strength of trophic competition within species (set to 1 in our simulations).
- *c*_*b*_ : the strength of trophic competition between species due to niche overlap between sympatric species (set to 0.3 in our simulations).
- *N*_*A*1_, *N*_*B*1_ : the number of individuals of species 1 with phenotype A and B respectively.
- *N*_*A*2_, *N*_*B*2_ : the number of individuals of species 2 with phenotype A and B respectively.
- *K* : the carrying capacity of the environment shared by the two sympatric species.
- *R*_*A*1_ : reproduction term describing the generation of individuals of phenotype A in species 1 through sexual reproduction.

The reproduction success *R*_*A*1_ counts the number of *A1* individuals produced across all possible mating combinations. Because heterospecific mating does not produce any offspring, *R*_*A*1_ decreases with the rate of reproductive interference, termed **I**. Because of the choice of an haploid model with one locus coding for colour pattern, only two mating combinations can result in the birth of *A*1 individuals (see fig. 1) :

- assortative mating : mating between two *A*1 parents, producing only offsprings with phenotype A
- disassortative mating : mating between an *A*1 individual and a *B*1 individual, resulting in 1/2 of offspring with phenotype A.

**Figure 1.**
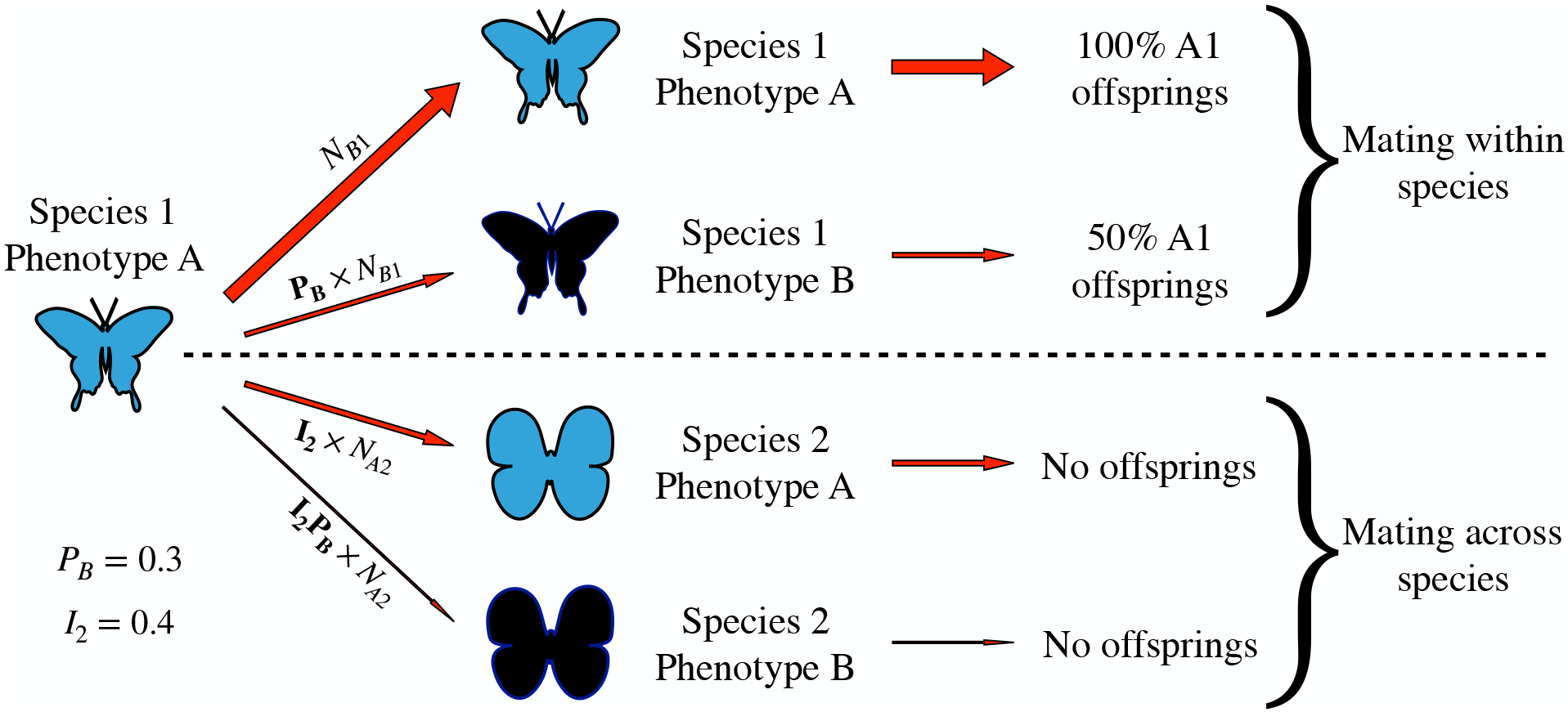
Mating behaviour of individuals of species 1 displaying the phenotype A and proportion of the different offsprings generated. Each individual of each species has four different possible partners. It can mate with a conspecific or heterospecific individual and for each of these two options, it can choose an individual displaying the same pattern (assortative mating) or not (disassortative mating). The frequency of each mating combination is determined by the values of **P** and **I**. Phenotypes A and B are represented by different colours (blue and black respectively) while species are represented by different shapes (species 1 is represented by long wings and a slender appearance and species 2 by short wings and a more bulky look). Each arrow corresponds to a given mating combination. The width of the arrow represents the number of times that combination occurs when assuming the parameters values shown on the bottom-left of the figure and equal population numbers : *N*_*A*1_ = *N*_*B*1_ = *N*_*A*2_ = *N*_*B*2_

The frequency of disassortative mating depends on individuals’ tendencies to depart from their assortative preferences, described by **P**. Following Kuno 1992 [19] and Kishi and Nakazawa 2013 [17], we consider that the frequency of each z mating combination depends on mate availabilities, related to their local densities *N*_*A*1_, *N*_*B*1_, *N*_*A*2_ and *N*_*B*2_. Hence, the reproduction term is written as :

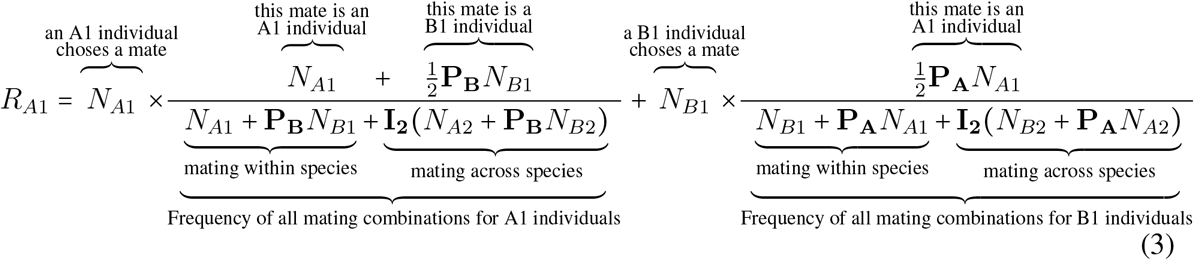

with **P**_**A**_ and **P**_**B**_ the tendencies to depart from assortative preference and reproduce with the alternative phenotype A and B respectively, and **I**_**2**_ the tendency of individuals from species 1 to engage in reproductive interference towards individuals from species 2. Given that assortative mating is widespread in species involved in Müllerian mimicry, we only explore departure from strict assortative mating ranging from **P** = 0 (individuals only mate with their own phenotype) to **P** = 1 (individuals mate at random regarding phenotype).

#### 3.3.2 Adult survival

Adult survival depends on the predation rate, which is modulated by mimicry (see supp. fig. S2 for more detail).

Following the model of Müllerian mimicry described in Joron and Iwasa 2005 [14], we assume that the predation risk on individuals with a given phenotype decreases with their density modulated by the individuals’ defence levels **L**. We assume that defence levels are equal within species but can differ between species, leading to asymmetrical contribution of the two mimetic species in predator learning when **L_1_** ≠ **L_2_**. The contribution of adult survival to the change in density of individuals with phenotype A in species 1, termed *S*_*A*1_, is described by :

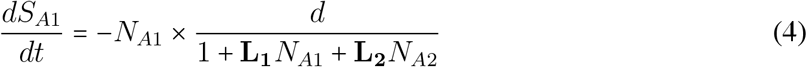

with

- *d* the intrinsic death rate, equal for all individuals regardless of species or phenotype,
- **L**_**1**_ and **L**_**2**_ the defence levels of individuals in species 1 and 2 respectively.

We use a number-dependent model of mimicry, as recommended in Mallet and Joron 1999 [23], because it results in a predation rate decreasing exponentially with prey population size and not linearly as in frequency-dependent models. This allows the number of preys killed by predators to converge towards a maximal threshold value when the prey population size increases (see supp. fig.S3). This threshold represents the number of prey that a predator community needs to consume before associating a mimetic pattern with defence mechanisms and depends on predator behavioural traits rather than on the prey population size, provided that encounter rate, and thus prey density, are high enough.

#### 3.3.3. Dynamics of phenotypes within species

The dynamic of both phenotypes in both species is simulated by the four different equations in supplementary figure S1 following the example given here for individuals of species 1 with phenotype A :

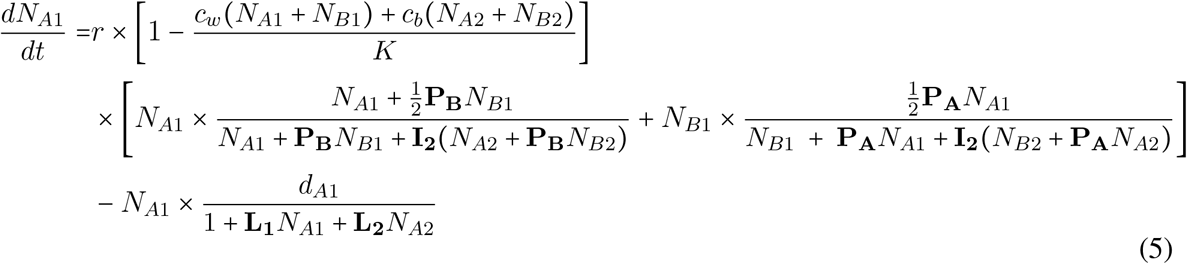

### 3.4 Numerical simulations

Because analytical resolution of this model is too challenging, we use numerical simulations with a wide variety of parameters values to explore the interactions between mimicry, assortative preference and reproductive interference (see table 1). We use a fixed level of trophic competition between species *c*_*b*_ = 0.3, lower than the trophic competition within species *c*_*w*_ = 1, and analyse this model in three steps.

**Table 1.** Default parameter values used in the simulations and range of values explored.

| Trait | Parameter | Values |
| --- | --- | --- |
| Intrinsic birth rate | $r$ | 0.2 for all individuals |
| Intrinsic death rate | $d$ | 0.1 for all individuals |
| Patch carrying capacity | $K$ | 1000 |
| Trophic competition |  |  |
| within species | $c_w$ | 1 |
| between species | $c_b$ | 0.3 |
| Defence mechanism strength |  |  |
| of species 1 | $L_1$ | $[0, 0.5]$ and $[0, 0.1]$ by steps of 0.001 |
| of species 2 | $L_2$ | $[0, 0.5]$ and $[0, 0.1]$ by steps of 0.001 |
| Tendency to mate with : |  |  |
| individuals with phenotype B | $P_B$ | $[0, 1]$ by steps of 0.1 |
| individuals with phenotype A | $P_A$ | $[0, 1]$ by steps of 0.1 |
| Tendency to attempt mating with : |  |  |
| species 2 | $I_2$ | $[0, 1]$ by steps of 0.1 |
| species 1 | $I_1$ | $[0, 1]$ by steps of 0.1 |

1. <u>Investigating the opposite effect of Müllerian mimicry and RI on colour pattern convergence</u>. First, we assume strict Müllerian mimicry where individuals from both species all have the same defence levels : **L**_**1**_ = **L**_**2**_. In this case, all individuals contribute equally to predator learning and only abundances determine which pattern offers better protection. We also assume individuals have the same tendencies to heterospecific mating : **I**_**1**_ = **I**_**2**_ and to disassortative mating : **P**_**A**_ = **P**_**B**_. We run simulations for values of *P*_*A*_ = *P*_*B*_ and *I*_1_ = *I*_2_ within the [0, 1] interval, using a 0.01 increment, and values of *L*_1_ = *L*_2_ within the [0, 0.05] interval, using a 0.001 increment, and we track down the evolution of the densities of each phenotype within each species until equilibrium is reached. We use uneven initial conditions with colour pattern polymorphism in both species and a slight disequilibrium in species densities and in colour pattern frequencies between species (*N*_*A*1_ = 50, *N*_*B*1_ = 60, *N*_*A*2_ = 70, *N*_*B*2_ = 40). These uneven initial conditions are based on previous smaller scale simulations and enable both colour pattern divergence and convergence between species, allowing to explore the interactions between the evolutionary forces studied.
2. <u>Investigating the effect of initial phenotype and species densities</u>. Because the evolutionary pressures studied are density-dependent, initial conditions can have a strong effect on the phenotype variations and species abundance at equilibrium. We thus investigate the impact of initial conditions on intra and inter-specific variations in colour pattern . We vary initial numbers of each phenotype within each species between 50 and 1000 by steps of 50 and assess the evolutionary scenario observed for each combination. To isolate the effect of initial densities from that of parameter values, we use two different sets of parameters values, one that led to convergence (**L**_**1**_ = **L**_**2**_ = 0.5, **P**_**A**_ = **P**_**B**_ = 0.8 and **I**_**2**_ = **I**_**2**_ = 0.3) and one that led to divergence (**L**_**1**_ = **L**_**2**_ = 0.5, **P**_**A**_ = **P**_**B**_ = 0.8 and **I_**2**_ = **I**_**2**_** = 0.8) in our previous simulations (see section 4.1). Both sets of parameters led to similar results when varying initial abundances (see fig.3 and supplementary figure S6). To describe the effect of initial species and phenotypic abundance, we use the following variables, computed with initial densities :

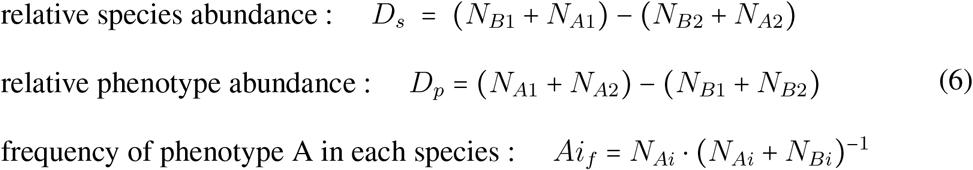

with *i* ∈ {1, 2}
3. <u>Investigating the effect of unbalanced evolutionary pressures between species..</u> Finally, we perform simulations assuming different levels of defence (**L**_**1**_ ≠ **L**_**2**_), reproductive interference (**I**_**2**_ ≠ **I**_**1**_) between the two species and assortative mating strength (**P**_**A**_ ≠ **P**_**B**_) between the two phenotypes. This allows for a more detailed study of a wider array of cases of mimicry and notably Batesian and quasi-Batesian mimicry. The two latter designate cases where :

- Batesian mimicry : one species shows no defence (either *L*_1_ = 0 or *L*_2_ = 0);
- quasi-Batesian mimicry or quasi-Müllerian mimicry : both species show strictly positive yet unequal level of defence (*L*_1_ ≠ *L*_2_, *L*_1_ and *L*_2_ > 0). We run simulations for values of *P*_*A*_ = *P*_*B*_ and *I*_1_ = *I*_2_ within the [0, 1] interval, using a 0.01 increment, and values of *L*_1_ = *L*_2_ within the [0, 0.01] interval, using a 0.001 increment.

We use the same unbalanced initial population numbers as in the previous simulations, also to highlight the effects of the three main evolutionary pressures. The default parameter values and ranges explored in the different simulations are detailed in tab 1.

Simulations are implemented in C++ using the Dormand–Prince (RKDP) method and a time-step of 1, compiled using the GCC g++ 4.2.1 compiler and run until population numbers vary by less than 10^−6^ over 500 consecutive units of time to ensure that equilibrium is reached. For each parameter set, the equilibrium density of each phenotype within each species is recorded. We analysed this data using the software R Studio 1.0.143 to determine the link between parameters’ values, <u>i.e</u> evolutionary pressure strength, and the evolutionary scenario observed. We define four main evolutionary outcomes :

- Convergence : both species display the same phenotype (either A or B) at equilibrium because the other phenotype has gone extinct.
- Divergence : the two species display a different phenotype at equilibrium because the alternative phenotype has gone extinct within each species.
- Polymorphism : polymorphism persist within at least one species at equilibrium. The other species persist with either a monomorphic or polymorphic state.
- Species exclusion : one species go extinct. The remaining species could then have either a single or both phenotypes maintained.

## 4 Results

We aimed at assessing the role of ecological interactions on the evolution of warning patterns in sympatric defended species. We explored the effects of six parameters involved in colour pattern variations within locality : the level of defence in species 1 and 2 (*L*_1_ and *L*_2_ respectively), the individuals tendency to depart from assortative mating and favour mating with the alternative phenotype (**P**_**A**_ and **P**_**B**_ respectively), as well as the tendency to heterospecific mating behaviour for species 1 and 2 (**I**_**2**_ and **I**_**1**_ respectively).

### 4.1 Reproductive interference limits convergent evolution of colour patterns in Müllerian mimics

First, we assume equal defence levels in both species (**L**_**1**_ = **L**_**2**_), therefore depicting conditions for Müllerian mimicry between species. We also assume equal species preference (**I**_**2**_ = **I**_**1**_) and phenotype preference (**P**_**A**_ = **P**_**B**_) in mate choice for all individuals. We then study the equilibria reached with different values of these parameters (Fig. 1).

#### Müllerian mimicry favours convergence

When individuals mate mostly within species (low values of *I*_1_ and *I*_2_) or when species mate at random regarding phenotype (**P**_**A**_ and **P**_**B**_ close to 1), individuals in each species face similar costs resulting from sexual interactions. In those cases, the evolutionary pressures due to sexual interactions have little impact on colour pattern variations. This results in a convergent evolution toward the phenotype A (blue area in Fig. 1 E and F), initially more common in the patch, as expected with a deterministic model of Müllerian mimicry without sexual interactions.

#### Assortative mating and RI promote colour pattern divergence

While convergence is widespread when individuals mate regardless of phenotype (**P**_**A**_ and **P**_**B**_ close to 1), increased assortative mating (**P**_**A**_ and **P**_**B**_ close to 0) tends to favour divergence (purple area in Fig. 1 A to D). Assortative mating indeed promotes the fixation of the most abundant phenotype within each species. Since the initially most common phenotype differs between the two species in our simulations, divergence is favoured. This divergence is however only observed if the costs associated with RI are high enough. Because the cost of RI is stronger between identical phenotypes across species (due to individuals’ tendency to assortative mating), it tends to favour colour pattern divergence between species. Indeed, divergence reduces the frequency of heterospecific mating behaviours since individuals tend to mate assortatively. Divergence is thus favoured, despite the strong impact of predation favouring convergence to phenotype *A* through Müllerian mimicry. Sexual behaviours thus interfere with evolutionary pressures stemming from prey-predator interations and can provoke colour pattern divergence in sympatric defended species, where convergence would be expected otherwise.

#### Müllerian mimicry prevents species exclusion

Species exclusion (yellow, dark blue, orange and red areas in fig. 2 C-F) is observed only when individuals have low levels of defences : **L**_**1**_ = **L**_**2**_ ≤ 0.005 (see Supplementary figure S5 for details). The higher predation rate generated by limited levels of defences combined with trophic and sexual competition leads to the exclusion of the initially least abundant species. Species exclusion also strongly depends on mate preferences tuning the strength of reproductive interference (see fig.S5). Both species persist when the levels of defence are high enough, highlighting the positive effect of Müllerian mimicry on species coexistence, despite competition.

**Figure 2.**
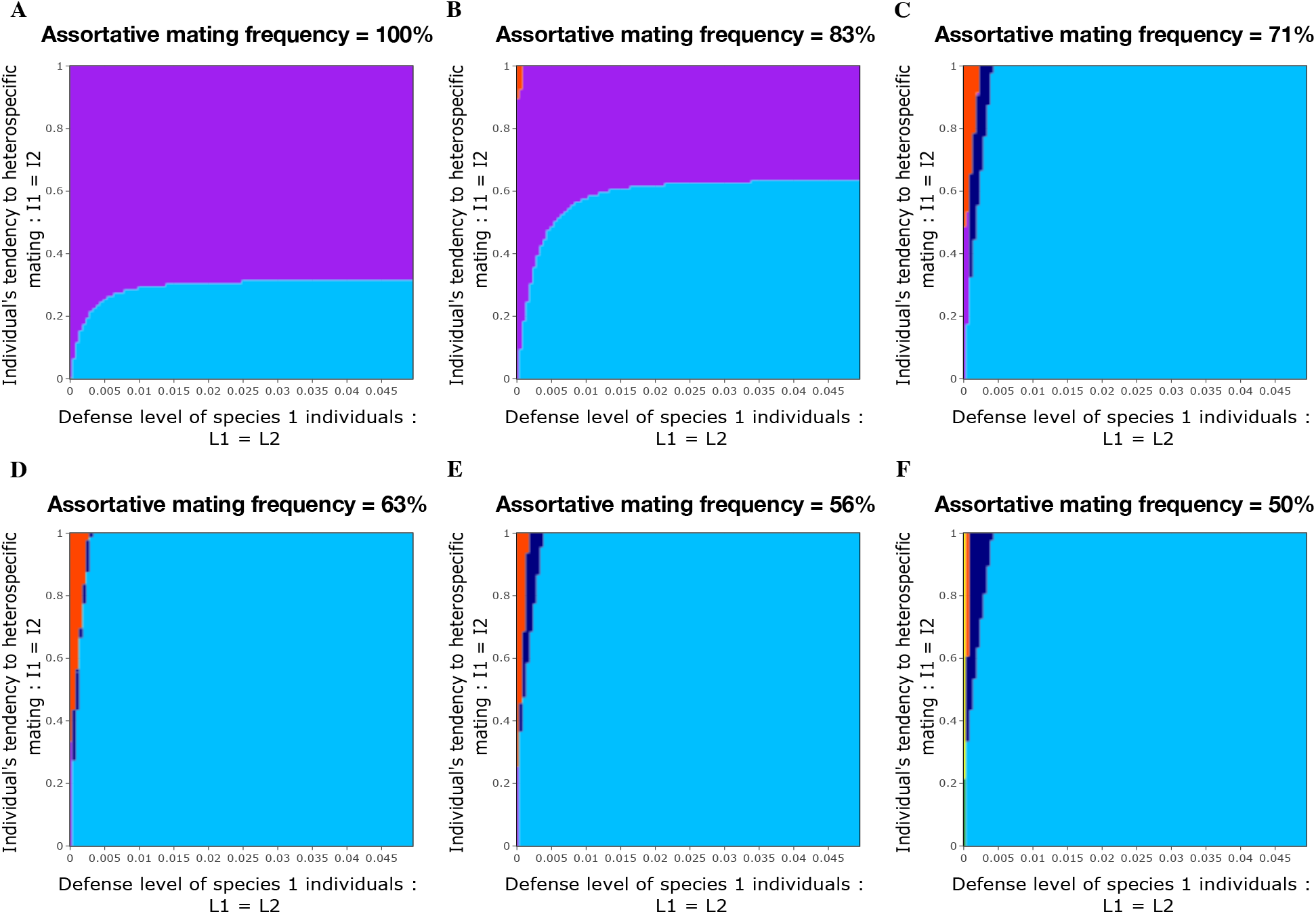
Effect of individual defence levels, RI costs and phenotype preferences on the warning pattern landscape at equilibrium between sympatric species. Each plot describes the equilibria obtained assuming a fixed phenotype preference equal for both phenotype (**P**_**A**_ = **P**_**B**_) and different values of defence trait (**L**_**1**_ = **L**_**2**_) and species preference (**I**_**2**_ = **I**_**1**_). Each colour represents an evolutionary outcome at equilibrium : Purple areas show the parameter sets leading to divergence between species and blue areas the ones leading to convergence toward the ph enotype A in both species. Dark blue areas and orange areas correspond to scenarios where only one phenotype and species persisted (*A*1 and *B*1 respectively). Scenarios where at least one species remains polymorphic occur when defence levels are low for both species and are shown in yellow and green colors (see Supplementary figure S4 for details).

#### Polymorphism results from the absence of the main evolutionary drivers

Polymorphism within both species is observed only in cases of random mating, without any mimicry and low RI costs (green area in the bottom-left corner of fig. 2F). The moderate trophic competition assumed here allows the persistence of both species and both phenotypes within each species.

### 4.2 Initial conditions strongly impact convergent evolution between sympatric species

#### Divergence is favoured when a different phenotype is initially more abundant in each species

As expected, when phenotype A frequencies are strongly unbalanced between the two species at initial state, colour pattern divergence is observed (pink and purple areas on fig. 3 A). The most frequent phenotype within each species becomes fixed, regardless of the difference in species abundances. Such divergence is more likely to occur when species densities are similar, as highlighted by simulations assuming different relative species densities (fig. 3B).

**Figure 3.**
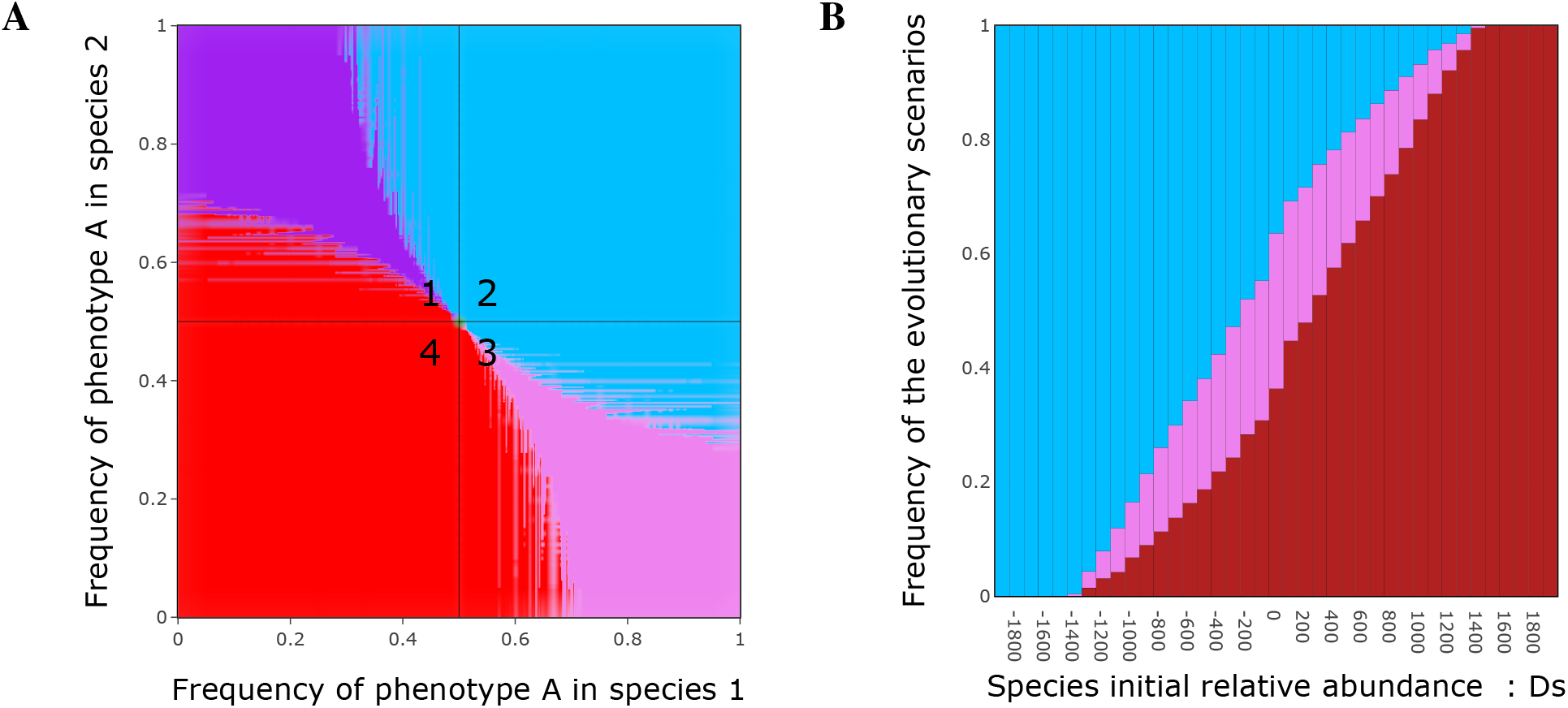
A. Effect of initial phenotype frequencies in each species on the convergence observed at equilibrium. B. Effect of relative species abundances on the frequency of each evolutionary scenario observed in simulations where, A is initially most abundant in species 1, and B in species 2. (*i.e.* as in the bottom-right square (3) on panel A). Each colour represents a different evolutionary scenario. Pink areas illustrate the divergence scenario where species 1 fixes phenotype A and species 2 fixes phenotype B. Purple areas illustrate the divergence scenario where species 1 fixes B and species 2 fixes A. Blue and red areas are associated with convergence towards phenotype A and phenotype B respectively. Note that the transparency levels on panel A indicates the frequency of the evolutionary scenarios observed in cases where several scenarios occurred for the same initial phenotypic frequencies, as a result of different initial species abundances, as detailed on panel B.

#### Convergence is favoured when the initially most common phenotype is shared between species

Convergence always occurs when the same phenotype is initially most abundant in both species (top-right (2) and bottom-left (4) squares on fig.3A). When a different phenotype is initially more abundant in each species however (top-left (1) and bottom-right (3) squares on fig.3A), both convergences and divergences occur depending on relative species abundance. Although convergence toward the phenotype most abundant overall initially is most frequently observed (result not shown here), it is not systematic, contrary to what can be expected in classical Müllerian settings. In those cases, reproductive interference makes both divergences or convergences possible (squares 1 and 3 in fig.3) and the outcome depends on relative species abundance, as illustrated in fig.3B.

#### Relative species abundance may change the direction of convergence

When a different phenotype is most abundant in each species initially, the relative species densities can result in both species converging toward the phenotype with the overall lowest initial density. Intriguingly, when species 1 has a higher initial density as compared to species 2 (high values of *D*_*s*_) and initially contains a majority of phenotype A, a convergence toward the phenotype B (with low initial density) can be observed (fig.3B). This counter intuitive result stems from the effect of higher competition for resource (*c*_*w*_) among individuals from the most abundant species. The more intense competition for food within species (*c*_*w*_) than between species (*c*_*b*_) has a strong negative impact on the most abundant species, and therefore on the phenotype most frequently displayed in this species, as illustrated when tracking down the densities of individuals from both species displaying the two phenotypes (fig.4A). The limited effect of intra-specific competition in the other species which is rarer favours the alternative phenotype. Combined with the increasing costs of reproductive interference as the initially rare phenotype becomes more abundant, it ultimately leads to the extinction of phenotype A and the convergence of both species toward the phenotype with the lowest initial frequency.

**Figure 4.**
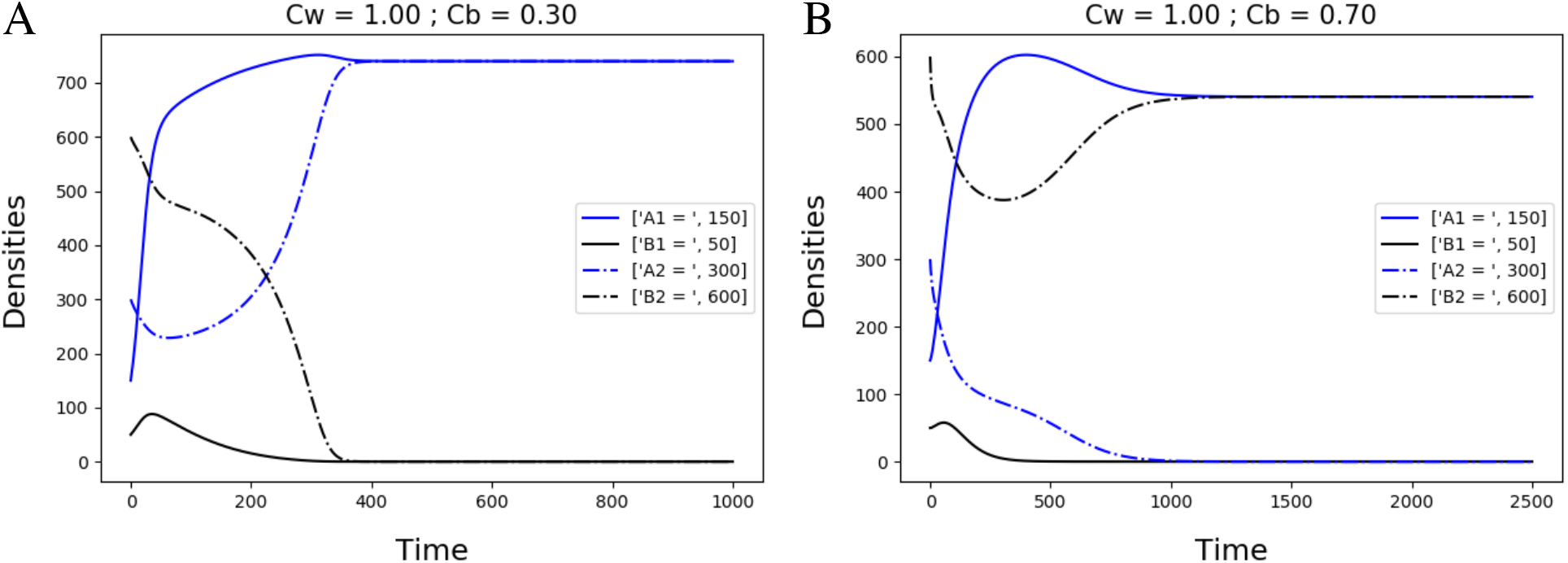
Direction of convergence shaped by trophic competition among species with contrasted initially density and phenotypic frequencies. The evolution of the densities of individuals from both species (plain and dotted lines for species 1 and 2 respectively) displaying either phenotype A or B (blue and black lines respectively) was tracked through time. Simulations were run assuming the same initially densities, with species 1 being the least abundant species and containing a majority of individuals with phenotype A, while the most abundant species, species 2, contained more individuals displaying the phenotype B. We compared the effect of trophic competition within and between species ; Panel A : *c*_*w*_ = 1 and *c*_*b*_ = 0.3 ; Panel B : *c*_*w*_ = 1 and *c*_*b*_ = 0.7

When trophic competition between species *c*_*b*_ increases with fixed competition within species *c*_*w*_, the advantages gained by individuals from the rarest species fade away and increased trophic and sexual competition lead to divergence (see fig.4B) or even the exclusion of the rarest species when *c*_*b*_ ≃ *c*_*w*_ (not shown here).

Competition for resources between mimetic species can thus strongly influence the direction of convergence, and therefore illustrates that the complex interactions between evolutionary pressures faced by mimetic species can drive phenotypic evolution in multiple directions.

### 4.3 Batesian mimicry enhances local convergence

Finally we explore the impact of unbalanced levels of defence between species and the interactions with mating behaviour and associated costs on the evolutionary convergence of colour pattern between sympatric species.

When the difference between the levels of defence of the two species increases, evolutionary divergence becomes less frequent (Fig. 5). In the Batesian or quasi-Batesian mimics, there is a strong positive selection on the phenotype most frequently displayed in the defended species, therefore promoting convergence. Divergence is unlikely because when it occurs, the undefended species loses the benefit from mimicking the species with defences whilst still enduring trophic and some reproductive costs associated with sympatry. Moreover, since the prey population of defended mimics is big enough, predator learning is barely impaired by the presence of undefended mimics meaning the pressure to diverge from these mimics is weak. Nevertheless, this convergence can be altered by differences in assortative mating strength and cost of reproductive interference. When mimicry benefits are balanced out by relative assortative mating and reproductive interference costs, the frequency of divergence increases. This is even more true since the most frequent phenotype is initially different in both species (Fig. 5A-B).

**Figure 5.**
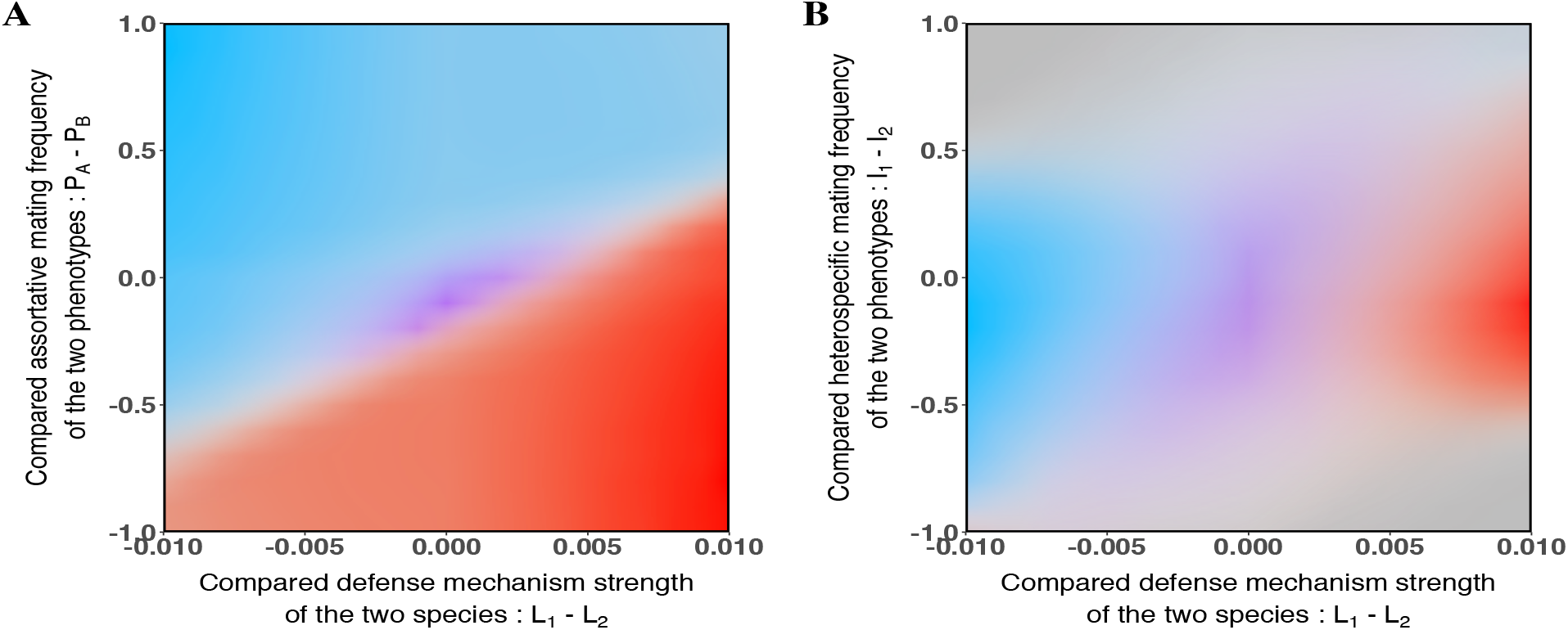
Convergence and divergence observed depending on species defence levels, phenotypic preferences (A) or costs of reproductive interference (B). For synthetic representation, we computed the differences between analogous parameters for the two species : *L*_1_ − *L*_2_, *I*_1_ − *I*_2_ respond to multiple couples of parameter values that result in the same difference. Colour intensity describe the frequency of each scenario in all the simulations with couples of parameters resulting in the same values of *L*_1_ − *L*_2_, *I*_1_ − *I*_2_ and *P*_*A*_ − *P*_*B*_. Each colour represents a different evolutionary outcome at equilibrium : blue is associated with convergence toward phenotype A, red is associated with convergence toward phenotype B and purple is associated with divergence where species 1 fixes phenotype B and species 2 fixes phenotype A.

## 5 Discussion

### 5.1 Mimicry enhances species co-existence despite reproductive costs associated with sympatry

Reproductive interference has been shown to frequently provoke species exclusion [17]. Nevertheless, our model revealed that by reducing predation pressure, mimicry among defended species significantly diminishes species extinction risk. Even for elevated cost of reproductive interference, this mutualistic interaction promotes species co-existence. This result is consistent with the persistence of large number of mimetic species in sympatry, as hihlighted in [1].

In completely undefended species, however, the cost of reproductive interference strongly enhances local risks of extinction when the sympatric species share the same mating cues. Density-dependent processes generated by either mutualistic interactions, such as Müllerian mimicry, or antagonistic interactions, such as reproductive interference, have a strong impact on species coexistence in sympatry, highlighting the eco-evolutionary feedback between the evolution of colour pattern within species and the extinction probabilities.

### 5.2 Mating behaviour interferes in the convergence of colour patterns between mimetic species

Our results suggest that assortative mating based on colour-pattern can increase phenotypic divergence between species provided that the most common phenotype is initially different within each species, that both species have similar levels of defence and all individuals are similarly assortative in their mating choice. By favouring the most frequent colour pattern within each species, assortative mating rapidly leads to the fixation of different colour pattern between species, even though Müllerian mimicry favours convergence. Assortative mating is frequently observed in species involved in Müllerian mimicry [13] and is expected to promote shifts in colour patterns in diverging species [2]. This is especially true if the postzygotic isolation between the sympatric species is strong [21], as assumed in our model. The evolution of assortative mate preferences is thus an important factor that may promote divergence in colour patterns between species, especially when the cost of reproductive interference is high for both diverging species. Indeed, high costs of reproductive interference for the two sympatric defended species can limit the convergent evolution of colour patterns. Multiple studies have already observed that sympatric species interacting through reproductive interference could coexist, as long as the costs faced by both species were balanced, and that divergence usually occurred [19, 18]. Field tests with mimetic species have also revealed that multiple mimicry rings can co-occur in spite of the convergent evolution promoted by Müllerian mimicry. Such diversity is possible if predator knowledge saturation is reached, meaning that predator can successfully recognise and avoid the different sympatric morphs which are therefore protected. This happens when all the signals are frequent enough for predators to memorise them all [5] or if the cost of predation on a particular conspicuous morph is so high that predators develop a general avoidance of all strongly conspicuous patterns [6]. Rare morphs in the population might also face lower predation rate because of predator neophobia [24] or when they exhibit classical warning colours [29] that predators generally tend to avoid. If one of the former mechanism allows a diversity of patterns to be maintained in the population, reproductive interference and mate selection might overcome the effect of mimicry, resulting in the diversification of warning patterns as observed in our model. To our knowledge however, little work has been done to investigate the potential conflict between mimicry and reproductive interference in spite of promising experimental results on the interaction between natural and sexual selection in mimicry rings [15, 6]. Our model thus highlights the need to investigate the costs associated with heterospecific sexual interactions to understand the evolution of divergent colour patterns within and between species, as well as the co-existence of multiple mimicry rings within locality.

### 5.3 Interactions between unbalanced defences and costs of RI lead to different scenarios of phenotypic diversification

Here, we observed that convergence in colour pattern might be favoured in cases of uneven levels of defence among sympatric species. This results is surprising because species involved in Batesian mimicry are frequently polymorphic [20] and Batesian mimicry is assumed to promote shift in colour patterns in the most defended species [8]. A key assumption of our model can explain these findings : the presence of undefended mimics does not impact the learning process of predators. This implies that co-existing with a less-defended mimic does not increase the predation risk suffered by individuals from the most defended species. Although a strong hypothesis, this assumption might actually be met in mimicry rings when the model species are highly defended or abundant enough so that the co-existence with a Batesian mimic does not have a significant negative impact on them. In that case, the strong advantage of mimicry promotes the evolution of the colour pattern displayed by the model in the least-defended species. Moreover, the evolutionary chase assumed to result from Batesian mimicry might not be as systematic as previously thought [15]. That is particularly true if the mimetic species exhibits some level of defence as in many of our simulations. In that case, high toxicity levels or frequent encounters with predators for the model species might be enough to create aversion for the pattern resulting in convergence across species.

### 5.4 Demographic history and niche overlap of sympatric species determine mimetic evolution at local scale

Our simulations investigating the effect of initial conditions on phenotypic evolution across species highlight that initial unbalance in phenotype frequencies across species promotes divergence, which is enabled by the presence of sexual interactions and notably reproductive interferences. Initial phenotype frequencies in the different species and initial species abundances are also shown to strongly drive the direction of convergence of warning patterns, especially when a different phenotype is most abundant in each species initially. This results from complex ecological interactions between tropic competition within and across species and mimicry. It occurs because the evolution of mimetic pattern relies on density-dependent processes generated by encountering rates, either (1) between predator and prey or (2) between mates or (3) between competitors. While positive density dependent selection exerted by predators may favour convergence of colour patterns between sympatric species, divergence of colour patterns can also be observed in species with similar abundances as a result of high heterospecific encountering rates and associated costs of RI. In particular, we show that the cost resulting from increased predation when maintaining two different patterns in the patch (divergence scenarios) is compensated by lower costs of sexual interactions due to less numerous heterospecific reproductive behaviours. The evolutionary history of colour patterns and of population sizes in sympatric species at local scale shapes the convergence or divergence of phenotypes within locality. Understanding the demographic history of populations of different species currently living in sympatry is thus crucial to understand the observed phenotypic convergences and divergences. Furthermore, by determining the heterospecific encountering rate and the level of trophic competition, niche overlap between mimetic species is likely to strongly shape the direction of selection on colour pattern. The evolution of mimetic coloration between species is frequently associated with convergence in other traits : in mimetic butterflies for instance, convergence in flight height [7] and host-plant [32] might result in enhanced reproductive interferences and trophic competition respectively. Variations in the levels of niche overlap and territoriality has been documented to influence the convergence vs. divergence of species recognition signals, driven by behavioral interferences between species [11]. The complex ecological interactions of mimetic species therefore interact with the stochasticity in population sizes and may result in contrasted evolutionary outcomes, that might explain the variations in mimicry rings observed within and across localities.

### 5.5 Conclusions & perspectives

Our model highlights that mate preference towards colour pattern and potential costs associated with heterospecific sexual interactions have a strong impact on phenotype evolution, even in the case of strong natural selection generated by Müllerian or Batesian mimicry, and may limit the convergent evolution of warning colorations in defended species. Hence for two species with high enough defence levels, divergence can be less costly than converging to the same phenotype because it reduces the reproductive interference costs that can emerge when convergence occurs. Our model also shows that surprising outcomes can occur as a result of complex interactions between evolutionary pressures, even when considering a deterministic evolution of population densities. Notably, species can end up converging to the initially least abundant phenotypes in the patch as a result of interactions between trophic competition, sexual competition and mimicry. Those interactions, especially reproductive interference, depend on the relatedness between species, because individuals from highly divergent species might display different trophic preferences and divergent behaviour toward other mating cues (such as odours) limiting confusion in mate recognition. The convergent evolution of colour pattern might thus be more impaired by cost of trophic competition and RI in closely-related species as compared to phylogenetically-distant ones. The effect of RI in the evolution of mimetic colour pattern might be especially important when investigating the phenotypic diversification in closely-related species, sharing similar ecological niches and suffering from high costs of RI. Empirical data on heterospecific sexual behaviour between closely related species involved in Müllerian mimicry are now requires to estimate the importance of RI and niche overlap in the evolution of mimetic colour patterns.

## Supporting information

All supplementary material

## Contributions

Grégoire Boussens-Dumon : Conceptualization, Methodology, Software, Formal analysis, Writing - Original Draft.

Violaine Llaurens : Conceptualization, Methodology, Writing - Original Draft, Project administration, Funding acquisition

## Acknowledgements

The authors would like to thank the whole ‘Evolution and Development of Phenotypic Variations’ as well as Charline Smadi, Ludovic Maisonneuve, Lea Terray, Thomas Aubier and Marianne Elias for feedbacks on the project.

## Funding

This work was funded by the Emergence(s) program from Paris city council to VL.

## Notes

### Competing Interest Statement

The authors have declared no competing interest.

