## Supplementary material for "Sex, Competition and Mimicry : an eco-evolutionary model reveals how ecological interactions shape the evolution of phenotypes in sympatry": All supplementary material

### S1 Supplementary material

#### S1.1 Complete model

$$
\left\{
\begin{aligned}
\frac{dN_{A1}}{dt} = \; & r_{A1}N_{A1} \times \left[1 - \frac{c_w(N_{A1}+N_{B1}) + c_b(N_{A2}+N_{B2})}{K_1}\right] \times \left[\frac{N_{A1} + 0.5\mathbf{P_B}N_{B1}}{N_{A1} + \mathbf{P_B}N_{B1} + \mathbf{I_1}(N_{A2} + \mathbf{P_B}N_{B2})}\right] \\
& + r_{B1}N_{B1} \times \left[1 - \frac{c_w(N_{B1}+N_{A1}) + c_b(N_{A2}+N_{B2})}{K_1}\right] \times \left[\frac{0.5\mathbf{P_A}N_{A1}}{N_{B1} + \mathbf{P_A}N_{A1} + \mathbf{I_1}(N_{B2} + \mathbf{P_A}N_{A2})}\right] \\
& - N_{A1} \times \frac{d_{A1}}{1 + \mathbf{L_1}N_{A1} + \mathbf{L_2}N_{A2}} \\
\frac{dN_{B1}}{dt} = \; & r_{B1}N_{B1} \times \left[1 - \frac{c_w(N_{B1}+N_{A1}) + c_b(N_{A2}+N_{B2})}{K_1}\right] \times \left[\frac{N_{B1} + 0.5\mathbf{P_A}N_{A1}}{N_{B1} + \mathbf{P_A}N_{A1} + \mathbf{I_1}(N_{B2} + \mathbf{P_A}N_{A2})}\right] \\
& + r_{A1}N_{A1} \times \left[1 - \frac{c_w(N_{A1}+N_{B1}) + c_b(N_{A2}+N_{B2})}{K_1}\right] \times \left[\frac{0.5\mathbf{P_B}N_{B1}}{N_{A1} + \mathbf{P_B}N_{B1} + \mathbf{I_1}(N_{A2} + \mathbf{P_B}N_{B2})}\right] \\
& - N_{B1} \times \frac{d_{B1}}{1 + \mathbf{L_1}N_{B1} + \mathbf{L_2}N_{B2}} \\
\frac{dN_{A2}}{dt} = \; & r_{A2}N_{A2} \times \left[1 - \frac{c_w(N_{A2}+N_{B2}) + c_b(N_{A1}+N_{B1})}{K_2}\right] \times \left[\frac{N_{A2} + 0.5N_{B2}\mathbf{P_B}}{N_{A2} + \mathbf{P_B}N_{B2} + \mathbf{I_2}(N_{A1} + \mathbf{P_B}N_{B1})}\right] \\
& + r_{B2}N_{B2} \times \left[1 - \frac{c_w(N_{B2}+N_{A2}) + c_b(N_{A1}+N_{B1})}{K_2}\right] \times \left[\frac{0.5\mathbf{P_A}N_{A2}}{N_{B2} + \mathbf{P_A}N_{A2} + \mathbf{I_2}(N_{B1} + \mathbf{P_A}N_{A1})}\right] \\
& - N_{A2} \times \frac{d_{A2}}{1 + \mathbf{L_1}N_{A1} + \mathbf{L_2}N_{A2}} \\
\frac{dN_{B2}}{dt} = \; & r_{B2}N_{B2} \times \left[1 - \frac{c_w(N_{B2}+N_{A2}) + c_b(N_{A1}+N_{B1})}{K_2}\right] \times \left[\frac{N_{B2} + 0.5\mathbf{P_A}N_{A2}}{N_{B2} + \mathbf{P_A}N_{A2} + \mathbf{I_2}(N_{B1} + \mathbf{P_A}N_{A1})}\right] \\
& + r_{A2}N_{A2} \times \left[1 - \frac{c_w(N_{A2}+N_{B2}) + c_b(N_{A1}+N_{B1})}{K_2}\right] \times \left[\frac{0.5N_{B2}\mathbf{P_B}}{N_{A2} + \mathbf{P_B}N_{B2} + \mathbf{I_2}(N_{A1} + \mathbf{P_B}N_{B1})}\right] \\
& - N_{B2} \times \frac{d_{B2}}{1 + \mathbf{L_1}N_{B1} + \mathbf{L_2}N_{B2}}
\end{aligned}
\right.
$$

Supplementary figure S1 – Complete model used to simulate the dynamics of two phenotypes (A and B) common to two different species (1 and 2). Each equation describes the temporal dynamic of one phenotype within one species. All individuals in the patch compete for food within and across species with intensity $c_w$ and $c_b$, depart from assortative mating with tendencies $\mathbf{P_A}$ or $\mathbf{P_B}$ depending on their own phenotype and from homospecific reproductive behaviours with tendencies $\mathbf{I_2}$ or $\mathbf{I_2}$ depending on their species. Death rates ($d_1$ and $d_2$), growth rates ($r_1$ and $r_2$) and costs related to competition for food ($c_w$ and $c_b$) are the same for all individuals.

#### S1.2 Predator behaviour shaping the evolution of mimicry in defended prey

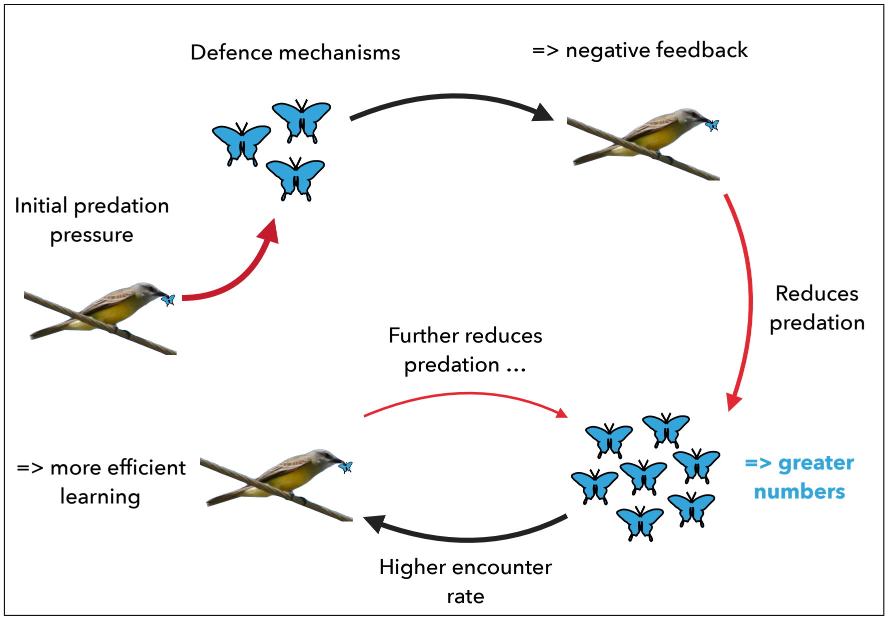


Supplementary figure S2 – Mechanism by which mimicry reduces predation pressure on mimetic defended individuals illustrated here with birds predating on blue butterflies. Prey defences limit predation due to the negative feedback for predators. Prey densities increase as a result which facilitates predator learning due to increased frequency of encounter with defended preys.

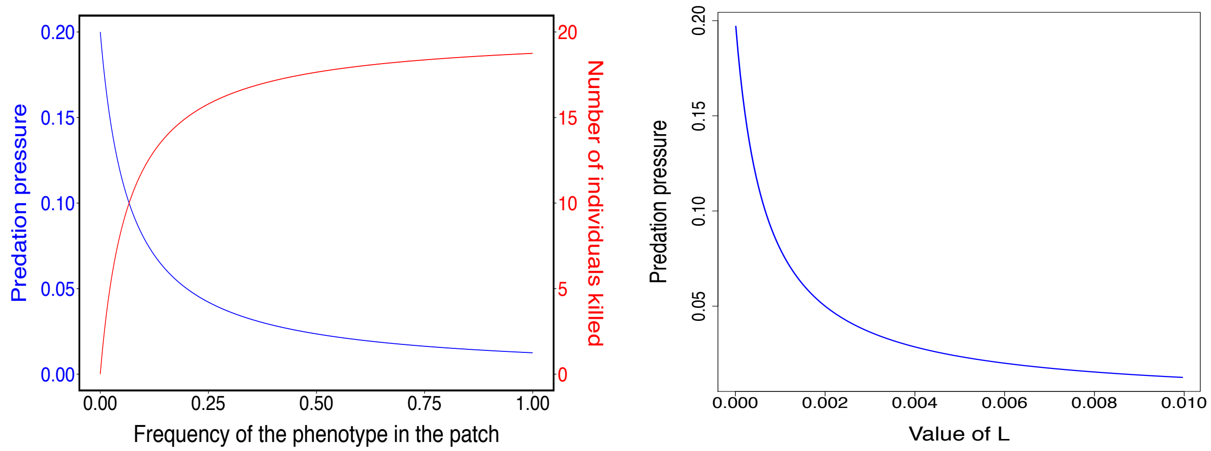


(a) Impact of the population size

(b) Impact of the defence mechanisms' strength

Supplementary figure S3 – Change in predation pressure relative to the prey population size and the strength of the defence mechanism, shown here for a single mimetic pattern. Fig. S3a illustrates the above mentioned exponential decrease of predation rate with population size. It also shows the convergence of the number of prey killed by predator towards a threshold representing the number of prey that predators need to consume before they associate mimetic pattern with defence mechanisms. Fig. S3b illustrates how the intensity of the defence mechanism reduces the predation pressure faced by mimetic individuals as stronger defence mechanisms are assumed to accelerate predator learning.

#### S1.3 Evolutionary scenarios when all individuals mate at random regarding phenotype

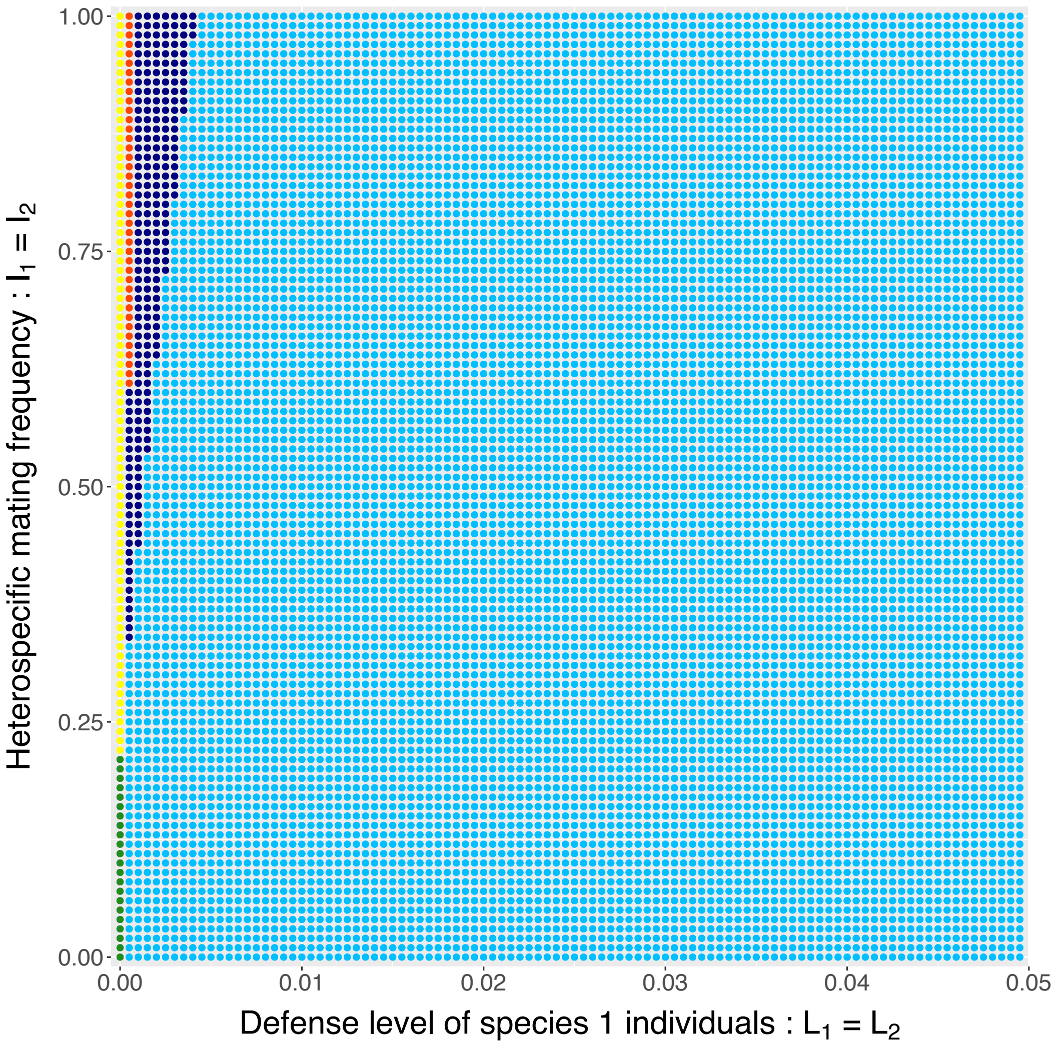


Supplementary figure S4 – Effect of the cost associated with reproductive interference and the level of defence of both species on the evolutionary scenario at equilibrium. When both species show high levels of defence, convergence to the initially most abundant phenotype occurs in spite of reproductive interference. When defence levels decrease however, the benefit of mimicry is balanced by the cost associated with reproductive interference resulting in divers scenarios of species exclusion (dark blue and orange areas). Finally, polymorphism is maintained in the absence of defence mechanisms in both species $L_1 = L_2 = 0$ as neither phenotype is favoured by those conditions. When reproductive interference costs are high, only one species persists (yellow area), and when they are low, both species remain (green area).

#### S1.4 Species extinction when one species shows no defence mechanisms : $\mathbf{L_1} = 0$ ou $\mathbf{L_2} = 0$

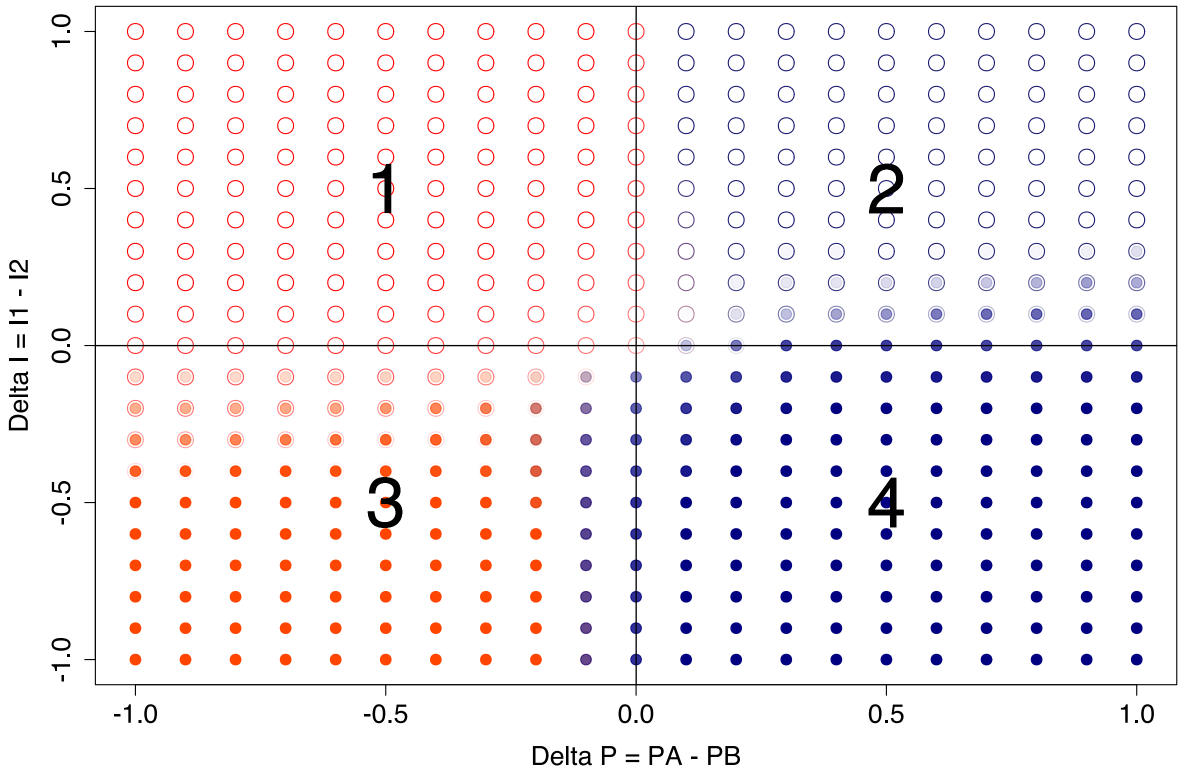


Supplementary figure S5 – Effect of the relative mating preference regarding phenotype (Delta P = $\mathbf{P_A} - \mathbf{P_B}$) and species (Delta I = $\mathbf{I_2} - \mathbf{I_1}$) on the phenotype remaining in case of species exclusion. Each colour represents one phenotype : dark blue for phenotype A and red-orange for phenotype B. Empty circles represent individuals of species 1 and full circles represent individuals of species 2.

When individuals show no defence trait, extinction becomes a frequent evolutionary scenario. Relative reproductive interference costs and departure from assortative mating then determine which morph remains within each species at equilibrium. This effect is stronger when initial population numbers are somewhat balanced, as in these simulations. High values of $\mathbf{P}$ and $\mathbf{I}$ mean individuals depart more frequently from assortative mating and homospecific mating respectively. Both result in a lower reproductive success (see S1). Therefore, squares 1 and 3 are associated with a lower reproductive success for phenotype A (higher values of $\mathbf{P_B}$) and squares 2 and 4 with a lower reproductive success for phenotype B (higher values of $\mathbf{P_A}$). Squares 1 and 2 are associated with a lower reproductive success for species 2 (higher values of $\mathbf{I_1}$) and squares 3 and 4 with a lower reproductive success for species 1 (higher values of $\mathbf{I_2}$). Because of their low reproductive success, the phenotype that departs the most from assortative mating and the species that faces the highest reproductive interference costs become extinct. Thus, only the individuals that mate most assortatively and within species remain at equilibrium.

#### S1.5 Evolutionary scenarios when varying initial conditions with parameter values resulting in divergence in the first simulations

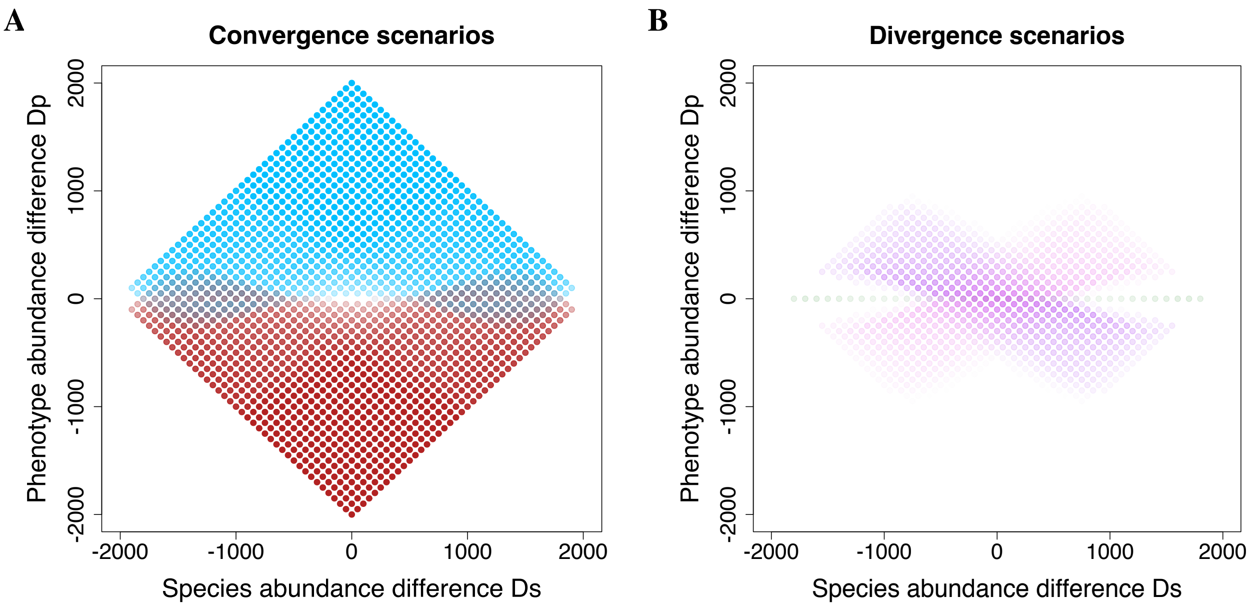


Supplementary figure S6 – Effect of initial relative species abundance $D_s$ and phenotype abundance across species $D_p$ on phenotypes at equilibrium when using parameter values that led to divergene in the first simulations ($L_1 = L_2 = 0.01$, $P_A = P_B = 0.2$, $I_1 = I_2 = 0.1$). Because each value of $D_s$ and $D_p$ can be associated with different values of initial abundances, different evolutionary scenarios can occur for any couple ($D_s$,$D_p$). The frequency of each scenario is represented by a level of opacity going from transparent (the scenario never occurs with the given values of $D_s$ and $D_p$) to fully opaque (the same scenario occurs systematically with the given values of $D_s$ and $D_p$). Each colour represents a different evolutionary scenario. Blue and red areas are associated with phenotype convergence across species toward phenotype A and B respectively. Pink areas illustrate the divergence scenario where species 1 fixes A and species 2 fixes B. Purple areas illustrate the divergence scenario where species 1 converges to phenotype B and species 2 to phenotype A. Finally dark green areas illustrate scenarios where A and B phenotypes remain at equilibrium in both species.
